# Engineered Fab glycosylation of a blood-brain barrier transporter single-chain variable fragment preserved functionality but did not impact anti-drug antibody formation

**DOI:** 10.1101/2025.11.28.691100

**Authors:** Gustaf Hederoth, Jielu Liu, Ana Godec, Nicole G. Metzendorf, Bence Rethi, Greta Hultqvist

## Abstract

The single-chain variable fragment (scFv) of the 8D3 antibody enables blood-brain barrier transport via transferrin receptor-mediated transcytosis. However, when fused to therapeutic antibodies, scFv8D3 increased immunogenicity and triggered anti-drug antibodies (ADA). Sialylation of N-linked glycans can critically influence protein immunogenicity, potentially conferring immunosuppressive properties and extended half-life. We investigated whether introducing sialic acid residues to scFv8D3 through engineered fragment antigen-binding (Fab) glycosylation could reduce ADA formation. We generated two novel variants with distinct N-linked glycosylation motifs (NXT/S), which both displayed sialic acid residues and retained functionality, as evidenced by the preservation of transferrin receptor binding and brain uptake. However, Fab glycosylation did not significantly alter pharmacokinetics or ADA production in mice. Our findings demonstrated that introducing Fab glycosylation is feasible, without compromising biological activity. Nonetheless, this strategy alone was insufficient to mitigate immunogenicity, underscoring the need for alternative approaches to improve the long-term safety of brain-targeting therapeutics.

**Statement of significance:** Sialylation of N-linked glycans can critically influence protein immunogenicity. We introduced N-linked Fab glycosylation sites into a scFv-based therapeutic construct with the aim of reducing ADA production upon repeated administration to mice. Although the sialylated variants retained functionality, pharmacokinetics and ADA levels remained unaffected, suggesting limited immunomodulation via engineered sialylation.

## Introduction

Biopharmaceuticals have greatly advanced the medical field, with monoclonal antibodies representing the largest class of newly FDA-approved drugs [1]. However, biologics are more likely to be recognized as foreign by the immune system, leading to anti-drug antibody (ADA) production. ADA can bind to the therapeutic, altering pharmacokinetics, and in some cases trigger hypersensitivity reactions or anaphylaxis.

While humanization has improved the safety of biologics, post-translational modifications can also affect drug immunogenicity. In particular, altered glycosylation and reduced sialylation of antibodies have been linked to inflammation and immune cell activation, as observed in rheumatoid arthritis [2,3]. Sialylated proteins can engage immunoregulatory receptors on immune cells, known as sialic acid-binding immunoglobulin-like lectins (Siglecs) [4]. Additionally, sialylated proteins can inhibit complement activation by recruiting Factor H, which promotes C3b cleavage [4,5].

A conserved N-linked glycosylation site is present in the IgG Fc region (N297) [6], however, over 10% of the circulating IgG antibodies also carry glycans in their fragment antigen-binding (Fab) regions [7]. Fab glycosylation increases markedly under certain pathological conditions, such as in rheumatoid arthritis or HIV-1 infection, where certain antibodies acquire Fab glycosylation sites through somatic hypermutations [8]. DNA segments convertible by single point mutations to encode NXT/S glycosylation motifs cluster notably at the complementary-determining regions of immunoglobulin variable genes suggesting that Fab glycans are evolutionarily conserved elements of antibody diversity [7]. Interestingly, Fab glycosylation has been shown to influence antibody-antigen affinity both positively and negatively, often exerting a relatively minor effect, which suggests a selection benefit not obviously explained by an increase in affinity [7,9,10]. Notably, when Fab glycosylation sites are inserted experimentally, i.e. not positively selected during B cell activation, the effect on antigen recognition may be more unpredictable.

The potential role of antibody glycans could be particularly interesting in therapeutic contexts. Intravenous immunoglobulin (IVIG), a preparation of pooled human IgG has been tested against a variety of disorders and is currently FDA-approved for 31 conditions [11]. IVIG possesses anti-inflammatory properties attributed to Fc sialylation of a fraction of the antibodies [12]. In mice, sialylated IgG acts by engaging SIGN-R1 (the murine homologue of human DC-SIGN) on splenic macrophages, leading to increased expression of the inhibitory receptor FcgRIIb [13]. Later evidence suggested that sialylation of the Fab region may contribute substantially to these immunomodulatory effects [14].

In this study, we analysed whether inserting N-linked glycosylation motifs into an antibody-based construct, and thereby increasing its sialylation, could confer immunosuppressive properties, reduce ADA production, and improve blood half-life. We studied the single-chain variable fragment of the 8D3 antibody (scFv8D3), which can facilitate blood-brain-barrier transport of large proteins via transferrin receptor (TfR) mediated transcytosis [15– 17]. Bispecific antibodies containing scFv8D3 elicit high ADA responses, limiting their therapeutic potential [18]. Sialylated and control scFv8D3 were injected into mice and we investigated their pharmacokinetics and ADA responses, with the overall goal of improving brain-targeting therapeutics.

## Materials and methods

### Design of glycosylated single-chain variable fragments

We studied the scFv8D3 structure in PyMOL and identified positions where single amino acid substitutions resulted in surface-exposed asparagine-X-threonine/serine (NXT/S) N-linked glycosylation motifs. Four favorable positions were selected: two near the complementarity-determining regions (D73N and S203N) and two more distal (L18T and K210N), and we generated a double-glycosylated (D73N/S203N) and a triple glycosylated (L18T/D73N/K210N) mutant. The corresponding genes, including an N-terminal Twin-Strep-tag, were synthesized and cloned into the pcDNA3.4 vector by GeneArt for recombinant expression (ThermoFisher).

### Expression and characterization of the scFv8D3 constructs

Proteins were transiently expressed in Expi293F cells and purified using Strep-Tag affinity chromatography [19]. The eluted proteins were concentrated using 30kDa centrifugal filters and buffer-exchanged to PBS. Endotoxin levels were confirmed to be below 0.09 EU per injection using a Limulus amoebocyte lysate assay (Microbial Analytics).

The purified proteins were analyzed using SDS-page as well as western blots using anti Strep-tag detection (Iba Lifesciences cat. no. 2-1507-001) and biotinylated Sambucus nigra agglutinin (Vector Laboratories cat. no. B-1305-2) for sialic acid detection. TfR binding of the scFv8D3 variants was measured by indirect ELISA. Plates were coated with recombinant murine TfR, blocked, and incubated with serial dilutions of the proteins. Binding was detected using an anti-Strep-tag antibody followed by HRP-conjugated anti-mouse IgG and development with TMB.

### Animal details

C57BL/6JBomTac mice (Taconic M&B), 3-4 months old, were kept in a controlled environment with free access to food and water. All experiments were conducted in accordance with Swedish ethical regulations and the European Directive 2010/63/E.U. on the protection of animals used for scientific purposes and were approved by the Uppsala County Animal Ethics Board (Ethical permit #5.8.18-04903-2022).

### Brain uptake and blood pharmacokinetics study

Mice received 1.5 μg (1.7 nmol/kg) of ^125^I-labelled scFv8D3 intravenously. For brain uptake, animals were perfused 6 hours post-injection; for pharmacokinetics, tail-vein blood samples were collected over 24 hours. Brains, organs, and tissues were harvested, weighed, and their radioactivity quantified using a gamma counter and expressed as percent injected dose per gram of tissue (%ID/g).

### Repeated injection of the scFv8D3 variants and evaluation of ADA responses

Mice received four intraperitoneal doses of 30 nmol/kg scFv8D3 variants at two-week intervals, followed by an intravenous tracer dose of ^125^I-labelled protein two weeks later. After 24 h, blood and tissues were collected to determine biodistribution and pharmacokinetics. ADA levels were measured in 1:100 diluted plasma samples by ELISA, using coated scFv8D3 (5 nM) and HRP-conjugated anti-mouse antibody as detection antibody.

## Results

### Characterization of N-linked glycosylated scFv8D3 variants

To assess the potential impact of glycosylation on the immunogenicity of an antibody fragment, we screened the variable regions of the 8D3 anti-TfR1 antibody for potential point mutations that introduce surface-exposed NXS/T glycosylation sites. We selected the D73N and S203N mutations to generate glycosylation sites near the CDRs and L18T and K210N at more distant sites. Since several naturally occurring antibodies carry multiple Fab glycans [8], we produced scFv8D3 mutant with either two (D73N/S203N) or three (L18T/D73N/K210N) glycosylation sites (Figure 1A).

**Figure 1.**
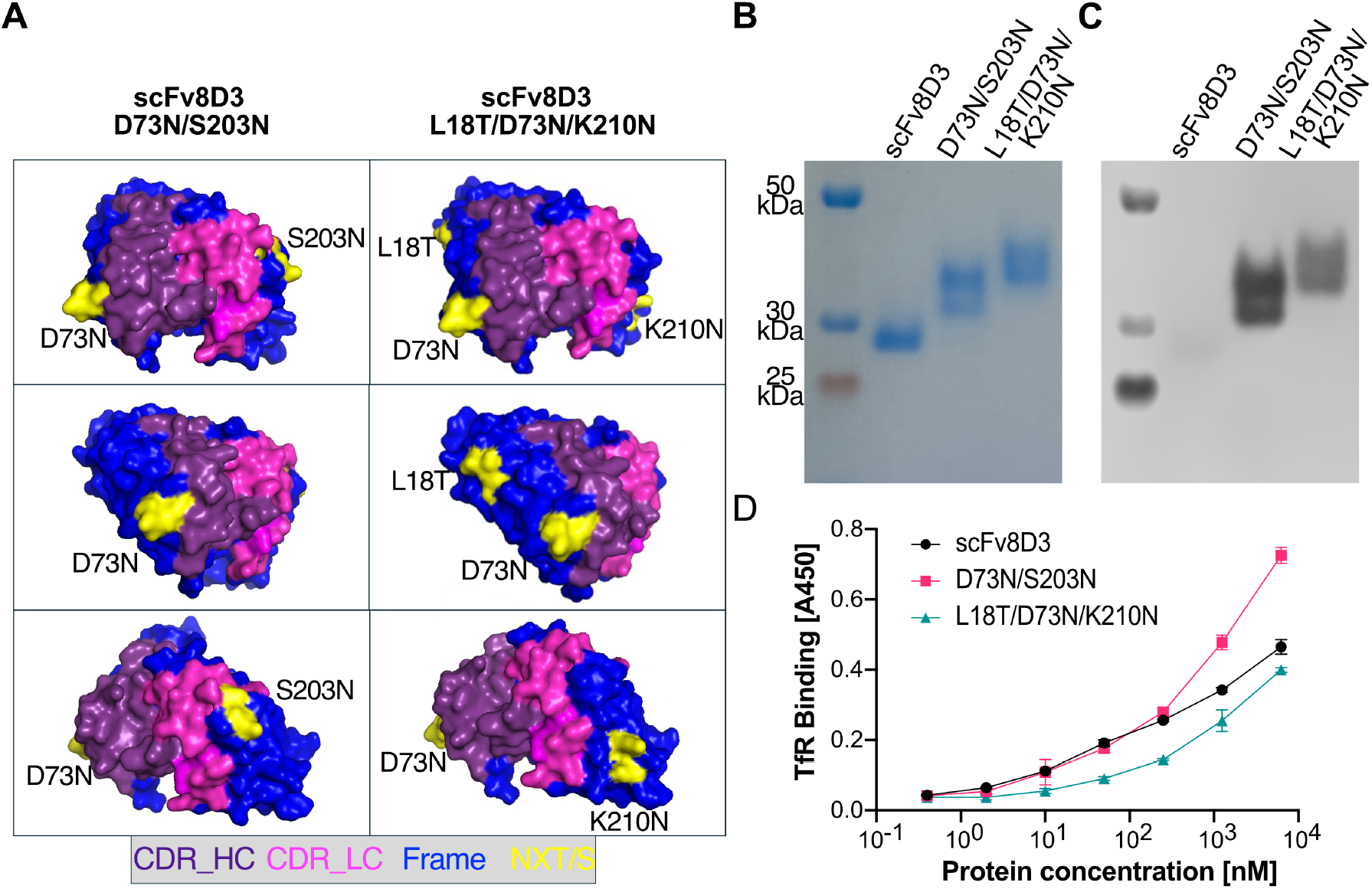
Design and characterization of glycosylated scFv8D3 variants. (A) AlphaFold 4 structure of scFv8D3 visualized in PyMOL, with complementarity-determining regions (CDR), framework regions, and positions of the inserted glycosylation motifs (NXT/S) highlighted. (B) SDS-PAGE analysis of Strep-tag purified scFv8D3 and glycosylated variants under non-reducing conditions. A pre-stained protein ladder was used to approximate molecular weights, which are predicted to be 29.6 kDa for all three variants based on the amino acid sequence. (C) Lectin blot with *Sambucus nigra* agglutinin visualizing the presence of sialic acid residues in the scFv8D3 variants. (D) Indirect ELISA demonstrating binding of the constructs to TfR. Data shown as mean ± SD.

The wild-type and glycosylated scFv8D3 variants were transiently expressed in Expi293F cells, purified via Strep-tag affinity chromatography, and analysed with SDS-PAGE and Western blot using an anti-Strep-tag antibody (Supplementary Figure S1). Introduction of the glycosylation motifs resulted in visibly higher molecular weights for the double- and triple-glycosylated mutants compared with the wild-type variant (Figure 1B), reflecting the added mass from post-translational modifications. The glycosylated and wildtype scFv8D3 variants exhibited similar thermal stability profiles (Supplementary Figure S1).

Sialic acid content was assessed by measuring *Sambucus nigra* agglutinin binding to the recombinant proteins [20]. As expected, wild-type scFv8D3, which lacks N-linked glycosylation motifs, showed no signal in the lectin blot, whereas both the double- and the triple-glycosylated mutants produced strong signals (Figure 1C). Interestingly, the scFv8D3 D73N/S203N mutant displayed a higher sialic acid signal compared with scFv8D3 L18T/D73N/K210N, suggesting more glycan incorporation but lower sialylation in the triple mutant.

As glycans near the CDRs could influence antigen binding, we assessed TfR recognition by the wild-type and the mutant scFv8D3 variants using ELISA. All three proteins exhibited robust antigen recognition, confirming that the added glycans did not disrupt the paratope (Figure 1D).

### Fab glycosylated scFv8d3 preserved brain targeting while modestly altering biodistribution and blood pharmacokinetics

To evaluate the impact of Fab glycosylation on pharmacokinetic properties, C57BL/6 mice were intravenously injected with 1.5 μg (1.7 nmol/kg body weight) of ^125^I-labelled scFv8D3 variants. Tissue distribution was analysed at 6- and 24-hours post-injection, and blood pharmacokinetics were assessed using sequential blood samples collected during the first 24 hours.

Wild-type and glycosylated scFv8d3 variants exhibited comparable brain uptake, although the brain levels of scFv8D3 D73N/S203N were slightly higher at 24 hours compared with the other constructs (Figure 2A). In plasma, the triple mutant L18T/D73N/K210N exhibited significantly higher levels at 6 hours compared with the other variants, while both glycosylated mutants showed a modest, non-significant increase at 24 hours relative to wildtype scFv8D3 (Figure 2B). Consistent with this, the estimated half-life of the glycosylated proteins was not significantly increased compared to the wildtype (Figure 2C). The glycosylated scFv8D3 variants also accumulated more in the liver, and the double mutant D73N/S203N showed higher levels in the spleen at 24 hours compared with the other constructs (Figure 2D-E). No major differences were observed among the proteins in other peripheral tissues and organs.

**Figure 2.**
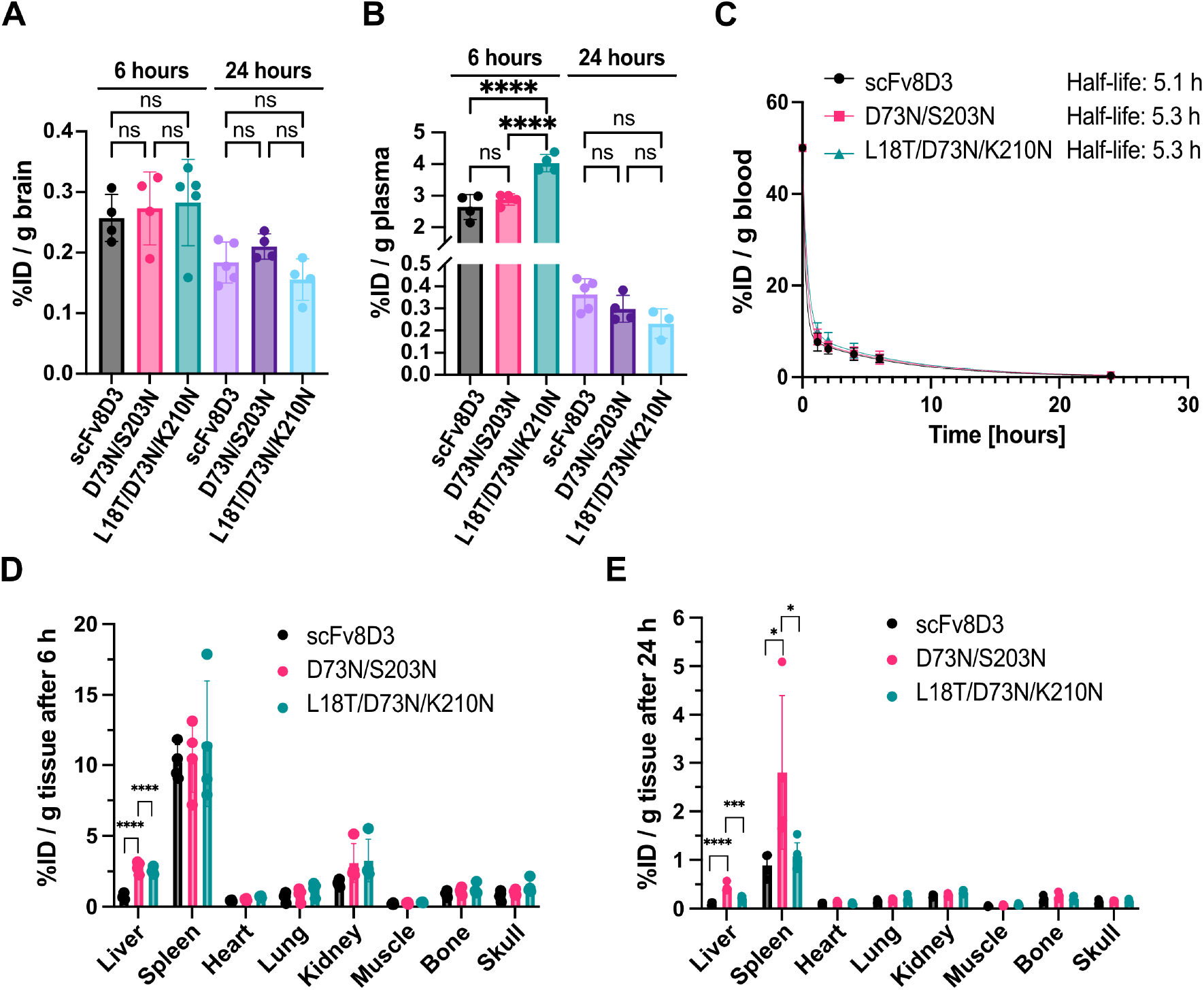
Effect of glycosylation on scFv8D3 pharmacokinetics and biodistribution. Brain uptake (A) and plasma levels (B) of radiolabelled scFv8D3 variants at 6- and 24-hours post-injection. (C) Blood pharmacokinetics of the scFv8D3 variants, determined from sequential blood samples and fitted using a two-phase decay non-linear regression model. (D,E) Biodistribution of the scFv8D3 variants 6 hours (D) and 24 hours (E) post-injection. Results are presented as mean±SD (n=4-5 animals per group). Tissue radioactivity is expressed as percent injected dose per gram of tissue (%ID/g). Statistical significance was assessed using one-way ANOVA followed by Tukey’s post hoc test for pairwise comparisons. P-values were defined as follows: ns > 0.05; * < 0.05; ** < 0.01; *** < 0.001; **** < 0.0001.

### Repeated dosing induced ADA irrespective of Fab glycosylation, which compromised scFv8d3 exposure and brain uptake

As ADA have been detected in response to TfR-binding antibodies [18], we investigated whether anti-inflammatory effects conferred by Fab glycosylation could modulate ADA production. Therapeutic doses (30 nmol/kg body weight) of scFv8D3 and the two glycosylated variants were administered to C57BL/6 mice using a sequential treatment regimen designed to induce ADA (Figure 3A). We assessed how repeated administration influenced the pharmacokinetic behaviour of the constructs. Evaluation of protein pharmacokinetics and half-life were complicated by the substantial variability observed between animals, even within the same treatment groups (Figure 3B). The biodistribution following repeated administration (Figure 3C) followed a pattern similar to that observed previously (Figure 2D-E). Notably, ADA production was observed in all three groups, regardless of the presence of glycosylation, although there was substantial variability within each group (Figure 3D).

**Figure 3.**
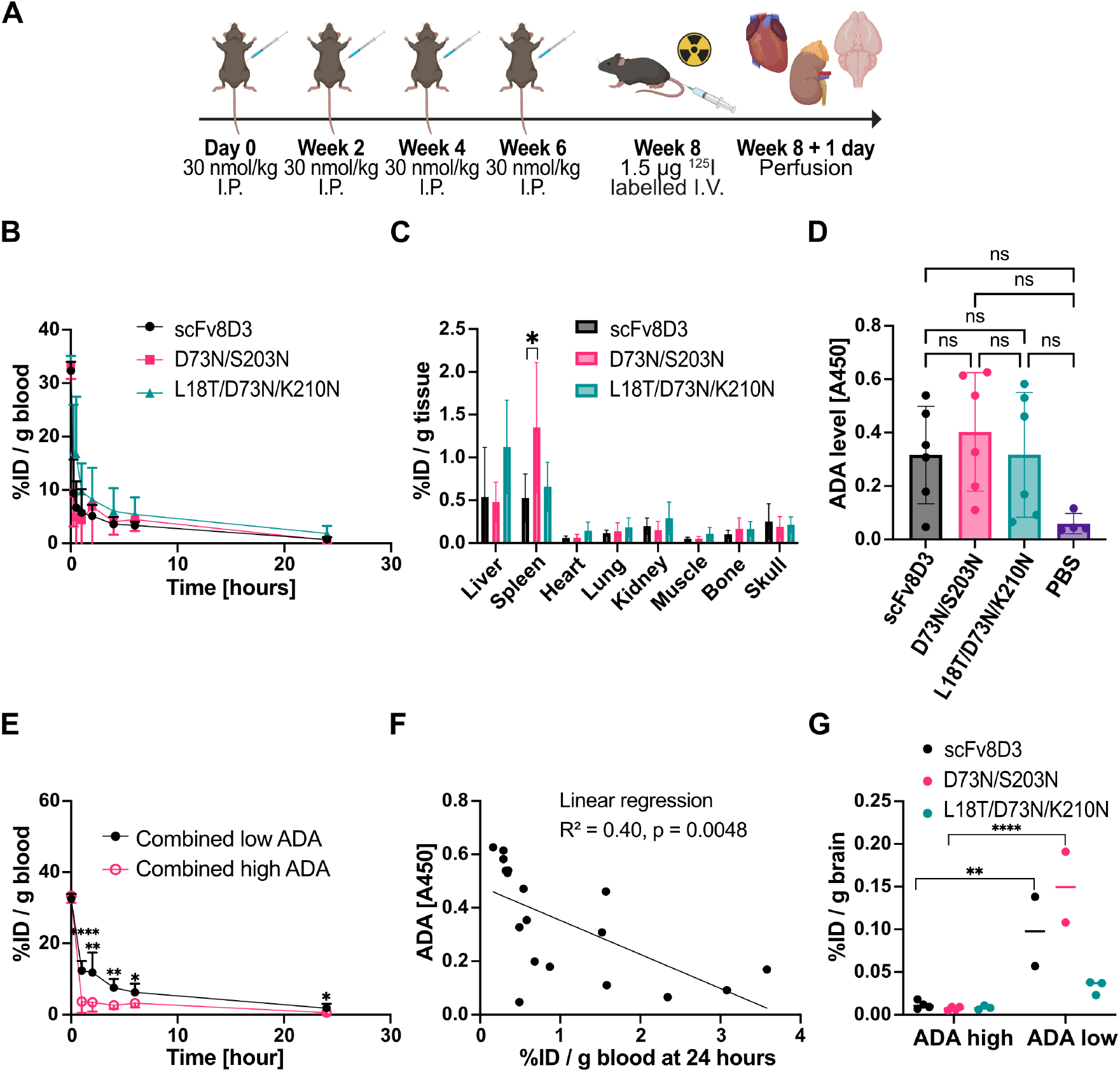
Repeated injections of non-glycosylated, double-, and triple-glycosylated scFv8d3 variants induce anti-drug antibodies (ADA) responses. (A) Experimental timeline showing repeated intraperitoneal (I.P.) dosing and a final intravenous (I.V.) administration of ^125^I-labelled scFv8d3. (B) Blood pharmacokinetics over 24 hours following the final injection. (C) Peripheral organ and tissue distribution. (D) ADA titers against the different scFv8D3 variants, with PBS-treated animals as a control group. (E) Impact of ADA status on blood pharmacokinetics. (F) Correlation between ADA titers and circulating scFv8D3 concentrations at 24 hours. (G) Brain uptake of scFv8D3 in animals classified by ADA levels. Statistical significance was assessed using one-way ANOVA followed by Tukey’s post hoc test for pairwise comparisons, except for Figure 3F, where Welch’s t-test was used. Data shown as mean ± SD (n=4-6 animals per group). P-values were defined as follows: ns > 0.05; * < 0.05; ** < 0.01; *** < 0.001; **** < 0.0001. Animals were classified as ADA-high or ADA-low based on an A_450_ value threshold of 0.2, corresponding to twice the highest value observed in PBS control animals (Figure 3D).

To assess whether the ADA response was responsible for the pronounced pharmacokinetic variability, animals were classified according to their ADA titers. Mice with high ADA levels exhibited consistently lower circulating concentrations of the scFv8D3 variants at all time points (Figure 3E). Consequently, total drug exposure, calculated from the area under the curve, was substantially reduced in these animals (Supplementary Figure S2). ADA titers negatively correlated with circulating scFv8D3 concentrations at the final time point, indicating that strong ADA responses accelerate protein clearance (Figure 3F). Moreover, high ADA titers were associated with a pronounced reduction in brain uptake, substantially compromising the brain-targeting function of scFv8D3 (Figure 3G).

## Discussion

Here we demonstrated that N-linked glycans can be introduced near the CDRs of scFv8D3 without disrupting antigen binding, as evidenced by successful transferrin receptor-mediated transcytosis of the modified variants (Figure 1D & 2A). The introduced glycans contained terminal sialic acids, confirmed by *Sambucus nigra* agglutinin binding (Figure 1C), which could potentially trigger immunosuppressive mechanisms [4]. Nevertheless, the glycosylated scFv8D3 variants displayed pharmacokinetic profiles comparable to the non-glycosylated protein, with no significant differences in blood half-life (Figure 2C). This contrasts with previous reports where sialylation through the addition of four glycosylation sites on erythropoietin and interferon alfa-2b resulted in a 3-fold and 25-fold increases in half-life, respectively [21].

Moreover, repeated injections of the scFv8D3 variants elicited comparable ADA levels, irrespective of the presence or absence of sialic acid (Figure 3D). We observed high ADA variability across all treatment groups, with higher ADA titers strongly associated with reduced scFv8D3 blood levels and markedly decreased brain uptake, effectively abolishing the antibody’s brain-targeting function (Figure 3E-G). Repeated scFv8D3 administration thus triggered ADA responses in mice that negatively affected protein availability and brain uptake, and that could not be reduced by the introduced sialylation.

The complete lack of effect of Fab glycosylation on ADA formation was somewhat unexpected, considering a key role of sialic acid in IVIG-mediated immunomodulation [11]. There is ongoing debate about the importance of the exact glycosylation sites in achieving anti-inflammatory effects [12,14,22] and such effects can also be context-dependent [23,24]. Our data nevertheless suggest that the induction of ADA responses may not be sensitive for anti-inflammatory signals elicited by sialic acid residues, which might be more effective in other contexts, such as regulating innate immune activation during inflammatory diseases [25].

## Acknowledgements

We thank Stina Syvänen and Dag Sehlin for their guidance in ADA studies. We acknowledge the Biophysical Screening and Characterization Unit at SciLifeLab for access to the Prometheus Panta instrument, as well as the Biophysical Screening and Characterization Radiochemistry Unit. Radiochemistry and animal work were performed at the SciLifeLab Pilot Facility for Preclinical PET-MRI, Uppsala University, Sweden, funded by the Knut and Alice Wallenberg Foundation.

