## Supplementary methods and supplementary figures for "Engineered Fab glycosylation of a blood-brain barrier transporter single-chain variable fragment preserved functionality but did not impact anti-drug antibody formation"

### Supplementary materials and methods

#### Recombinant expression and purification of scFv8D3 constructs

The proteins were recombinantly expressed in Expi293F cells (ThermoFisher cat. no. A14527) by transient transfection with the pcDNA3.4 vectors in the presence of polyethyleneimine (Polyscience cat. no. 24765–1) (22). The scfv8D3 variants were purified using a Strep-Tactin®XT 4Flow® FPLC column (Iba Lifesciences cat. no. 2-5024-001) according to the manufacturer's instructions. The eluted proteins were concentrated using Amicon Ultra 30kDa centrifugal filters (Sigma cat. no. UFC9010) and buffer-exchanged into PBS (ThermoFisher cat. no. 89882). Endotoxin levels were confirmed to be below 0.09 EU per injection using a Limulus amoebocyte lysate assay (Microbial Analytics).

#### Analyses of the expressed proteins

A total of 1 µg protein mixed with LDS sample buffer (ThermoFisher cat. no. B0007) was loaded onto a 4-12% Bis-Tris protein gel (Invitrogen cat. no. NW04125BOX). Electrophoresis was performed at 80 V for 90 minutes in MES buffer (ThermoFisher cat. no. NP0002). Gels were washed three times with deionized water and stained with PAGE blue protein solution (ThermoFisher cat. no. 24620). The gels were completely destained in deionized water and imaged using a digital camera. The PageRuler Plus Prestained protein ladder (ThermoFisher cat. no. 26619) was used to estimate molecular weight. To confirm identity of the expressed proteins the samples were transferred to a methanol activated PVDF membrane (ThermoFisher cat. no. 88520). After transfer, membranes were dried, reactivated, and blocked in 5% (w/v) skimmed milk TBS-Tween-20 overnight. The membrane was incubated with a mouse-anti Strep-tag antibody (Iba Lifesciences cat. no. 2-1507-001), washed thrice with TBS-Tween-20, followed by incubation with goat-anti-mouse conjugated to HRP (Sigma, cat. no. 12-349). The membrane was washed thrice with TBS-Tween-20. HRP chemiluminescent substrate (Invitrogen cat. no. WP20005) was added and imaged using the Odyssey Fc (LI-COR Biosciences).

#### Protein stability

Thermal unfolding of the scFv8D3 variants were measured using the Prometheus Panta (Nanotemper). Proteins were loaded into glass capillaries and heated from 25 °C to 80 °C with a linear temperature gradient of 1 °C per minute. The fluorescence intensities at 350 and 330 nm were monitored and the ratio was used to calculate the first derivative. The inflection temperature corresponds to a peak in the first derivative ratio which indicates a major protein unfolding event.

#### Determination of sialic acid presence

After electrophoresis as described above but using 0.5 µg of protein, samples were transferred to a methanol-activated PVDF membrane (ThermoFisher cat. no. 88520). The membranes were dried, reactivated, and blocked overnight in Carbo-Free Blocking Solution (Vector Laboratories cat. no. SP-5040-125). Biotinylated *Sambucus nigra* lectin (Vector Laboratories cat. no. B-1305-2) was added at a final concentration of 2 µg/ml with gentle shaking for 1 hour. Following three washes with TBS-Tween-20, membranes were incubated with avidin conjugated to horseradish peroxidase (HRP) (BioLegend, cat. no. 405103) at a final concentration of 0.6µg/ml for 45 minutes with gentle shaking. After three washes in TBS-Tween-20, HRP chemiluminescent substrate (Invitrogen cat. no. WP20005) was applied and imaged using the Odyssey Fc (LI-COR Biosciences).

#### Transferrin receptor binding

Binding to the murine transferrin receptor (TfR) was assessed using an indirect ELISA. High-binding 96-well plates (Sarstedt cat. no. 82.1581.200) were coated overnight at 4 °C with recombinant murine TfR extracellular domain (produced in-house) diluted in PBS to a final concentration of 2 µg/ml. The wells were blocked for 2 hours at room temperature with 1% BSA (VWR, cat. no. 421501J) in PBS while shaking at 500 rpm. Triplicates serial dilutions of the scFv8D3 constructs were added and incubated for 2 hours at room temperature with shaking. For detection, a mouse anti-Strep-tag antibody (Iba Lifesciences cat. no. 2-1507-001) was applied for 1 hour, followed by a goat anti-mouse antibody conjugated to HRP (Sigma cat. no. 12-349) for 45 minutes. K-blue aqueous TMB (Neogen Corp cat. no. 331177) was used for signal development, the reaction was stopped with 1M sulfuric acid (Thermo cat. no. 10794371), and absorbance at 450 nm was measured on a Spark multimode microplate reader (Tecan). All scFv8D3 variants and ELISA antibodies were diluted in PBS with 0.1% BSA and 0.05% Tween-20 (Sigma cat. no. P9416). The wells were washed with PBS containing 0.05% Tween-20 between every incubation step following the blocking step.

#### Animal details

C57BL/6JBomTac mice (Taconic M&B), 3-4 months old, were housed in the animal facility at Uppsala University. Animals were kept in a controlled environment (humidity: 50-55%, temperature 22-23 °C, 12:12-hour light:dark cycle) with free access to food and water. All experiments were conducted in accordance with Swedish ethical regulations and the European Directive 2010/63/E.U. on the protection of animals used for scientific purposes and were approved by the Uppsala County Animal Ethics Board (Ethical permit #5.8.18-04903-2022).

#### Radiolabeling of scFv8D3 variants

The scFv8D3 variants were radioactively labelled with iodine-125 ($^{125}$I) using the chloramine-T method. The proteins were mixed with PBS and $^{125}$I (Perkin Elmer Inc), after which 1 mg/ml of chloramine T (Sigma cat. no. 857319) was added to initiate the reaction. After 90 seconds, the reaction was quenched with 1 mg/ml of sodium metabisulfite (Sigma cat. no. 08982). Unbound $^{125}$I was removed by buffer exchange into PBS using spin columns (ThermoFisher cat. no. 89882).

#### Brain uptake and blood pharmacokinetics study

Brain uptake and biodistribution were evaluated in groups of four mice (both males and females evenly distributed), while blood pharmacokinetics were assessed in groups of five male mice. Each mouse received 1.5 µg (approximately 1.7 nmol/kg body weight) of $^{125}I$-labelled scFv8D3 via intravenous tail-vein injection. Experimental groups were randomly distributed among the cages. For the brain uptake study, mice were anesthetized with 3% isoflurane 6 hours post-injection and euthanized by transcardial perfusion with 0.9% (w/v) saline. For the blood pharmacokinetic study, mice were sampled from the tail vein at 1, 2-, 4-, 6-, and 24-hours post-injection using 8 µl capillaries (Vitrex Medical cat. no. 172613). Plasma was isolated from terminal blood after centrifugation. Perfused brains, peripheral organs (liver, spleen, heart, lung, kidney, thyroid), and tissues (muscle, bone, skull) were collected and weighed. Radioactivity was measured with a Wizard 1480 gamma counter (PerkinElmer) and expressed as percent injected dose per gram of tissue (%ID/g).

#### Repeated injection of the scFv8D3 variants and evaluation of ADA responses

Anti-drug antibodies (ADA) against the scFv8D3 variants were assessed in mice that received intraperitoneal injections of 30 nmol/kg of one of the scFv8D3 variants every two weeks, for a total of four doses. Two weeks after the final intraperitoneal injection, a tracer dose of 1.5 µg (approximately 1.7 nmol/kg) of $^{125}I$ -labelled scFv8D3 variant was administered intravenously. Blood, organs, and tissue were collected after 24 hours and measured as described above to determine %ID/g and evaluate blood pharmacokinetics.

ADA reactivity in plasma was measured by ELISA. High-binding 96-well plates were coated overnight at 4 °C with 5 nM of the respective scFv8D3 variant in PBS. The wells were blocked for 2 hours at room temperature with 1% BSA in PBS while shaking at 500 rpm. Plasma samples were diluted 1:100 and incubated overnight at 4 °C. For detection, goat anti-mouse antibody conjugated to HRP (Sigma cat. no. 12-349) was added for 1 hour at room temperature with shaking at 500 rpm. The assay was developed, and samples were diluted and washed as described above.

### Supplementary Figures

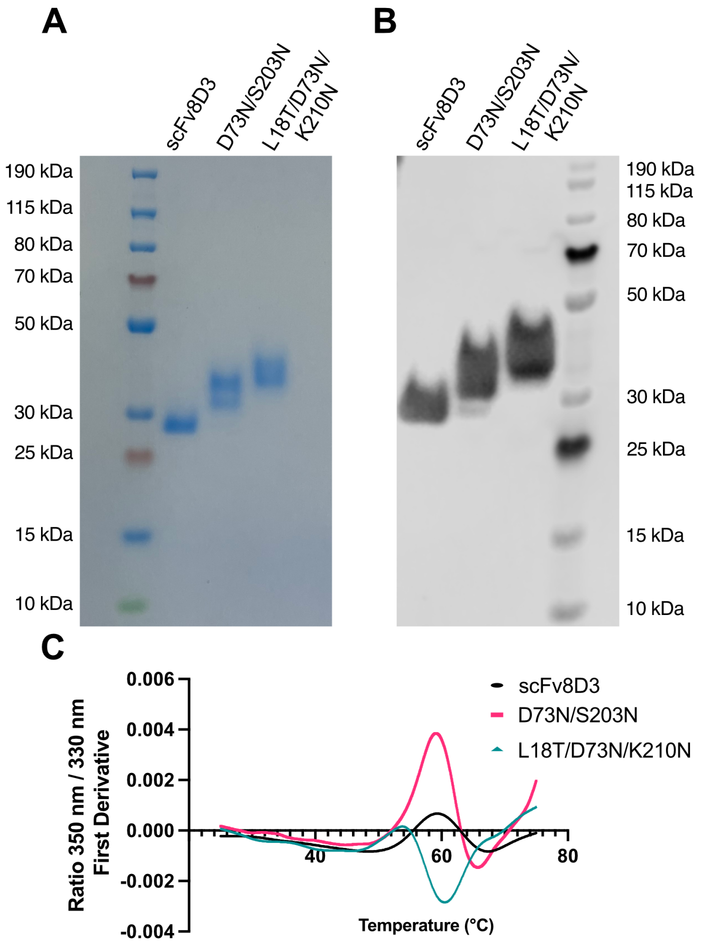


**Figure S1.** Characterization of protein purity, identity, and stability. (A) Full non-reducing SDS-PAGE gel analysis of Strep-tag purified scFv8D3 and scFv8D3 variants containing point mutations for N-linked glycosylation. A pre-stained protein ladder was used to approximate molecular weights, which are predicted to be 29.6 kDa for all three variants based on the amino acid sequence. (B) Western blot was utilized with an anti-Streptag antibody to validate the purified protein for variants with and without glycosylation sites. (C) Thermal unfolding Prometheus Panta data of the scFv8D3 variants. The inflection points are 59 °C for the wild-type scFv8D3. The scFv8D3 D73N, S203N has inflection points at 59 °C and 66 °C and the L18T, D73N, K210N has its inflection points at 54 °C and at 60 °C.

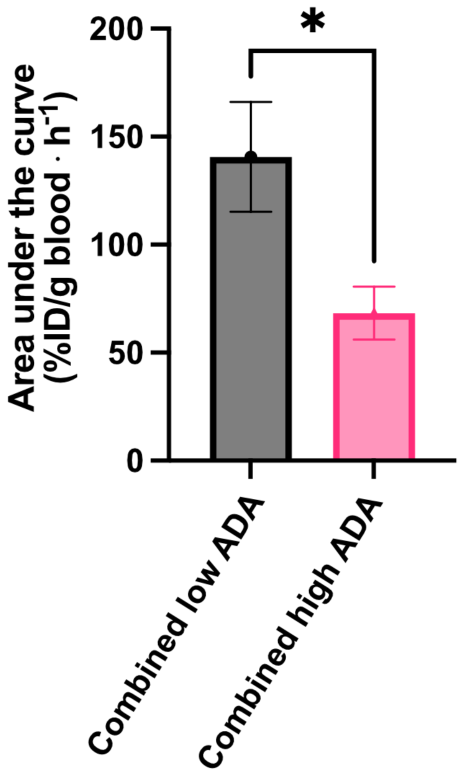


**Figure S2.** Total drug exposure for all three scFv8D3 variants combined based on ADA level. Area under the curve was calculated using the trapezoidal method on the blood pharmacokinetic curve in Figure 3E**.** Statistical significance was assessed using Welch's t-test. Data shown as mean ± SD. P-values were defined as follows: * < 0.05.
